# Extending the MAD Toolbox: New Polymer Builder and Enhanced Martini Database

**DOI:** 10.64898/2026.01.23.700524

**Authors:** Romuald Marin, Cécile Hilpert, Fabian Grünewald, Mariana Valério, Luís Borges-Araúj, Stéphane Janczarski, Nicolas O. Rossini, Siewert J. Marrink, Paulo C. T. Souza, Guillaume Launay

## Abstract

The MArtini Database (MAD) web server (https://mad.ens-lyon.fr/) has been updated with new tools, models, and capabilities to support a broader range of molecular systems for the Martini coarse-grained (CG) force field. The most notable addition is the MAD:Polymer Builder, enabling the automated construction of CG polymer models with varying architectures and complexities. The server now incorporates the latest developments in Martini 3, providing enhanced control within the MAD:Molecule Builder over the conversion of all-atom structures into CG representations, including the implementation of GōMartini 3 for protein complexes and water–protein interaction biases. The MAD:Polymer Builder and MAD:Molecule Builder are both adapted to work with intrinsically disordered proteins and domains. Substantial progress has been made in expanding the MAD:Database, making a growing library of Martini-ready compounds readily accessible across the entire MAD ecosystem. These advances position MAD as a comprehensive and evolving platform for the preparation of diverse systems in CG molecular simulations.

## Introduction

The Martini coarse-grained (CG) force field has become a widely adopted model for the molecular dyanamics simulation of biomolecules, soft matter, and complex materials systems due to its balance between computational efficiency and chemical specificity.^1^ Its continued development — most notably with the introduction of Martini 3^2^ — has extended its applicability to a broader range of chemistries, topologies, and interaction types, opening new possibilities for simulating increasingly complex systems at mesoscopic scales. A number of tools have been developed to facilitate the construction of complex molecular systems for the Martini 3 force field, such as bentopy,^3^ martinize2,^4^ COBY,^5^ TS2CG,^6,7^ polyply^8^ and others, each addressing different aspects of system assembly.

To support and streamline the modeling process for Martini users, we previously developed the MArtini Database (MAD) server,^9^ an online platform that facilitates the generation, conversion, and assembly of CG systems through modular tools and a curated database of preparameterized Martini models. Since its initial release, MAD has become an integral part of the Martini ecosystem, facilitating the use of tools such as martinize2^10^ and insane,^11^ and is now increasingly adopted by the CG community to construct Martini-compatible inputs for molecular dynamics (MD) simulations.

As the Martini force field has evolved, so too have the needs of its users. Growing demand for automated polymer construction, hybrid models, and more customizable workflows motivated a comprehensive update of the MAD platform. In this work, we present an extended version of the MAD server, hosted at a new location (https://mad.ens-lyon.fr), with major enhancements aimed at improving usability, interoperability, and modeling flexibility. The most significant addition is the MAD:Polymer Builder, based on the polyply framework,^8^ which allows users to build Martini-compatible CG polymer models from arbitrary sequences and architectures. In addition to synthetic polymers, it also supports carbohydrates^12,13^ and the generation of disordered peptides using the Martini 3-IDP extension,^14^ broadening its application to biomolecules. The Molecule Builder has also been expanded to incorporate support for GōMartini 3,^15^ enabling hybrid Martini/Gō-like modeling of structured dimers and oligomers, but also adding water biases for improving the hehavior of peptides^16,17^ and intrinsically disorder proteins.^18,19^ In parallel, the MAD:Database has been substantially enriched with new compounds, particularly the recently released Martini 3 lipidome^20^ and new lipid nanoparticle components.^21^ The codebase has also undergone a complete revision to improve performance, modularity, and future extensibility.

Together, these advances mark a substantial step forward in the capabilities of the MAD server, offering a unified, extensible, and user-friendly environment for building a wide variety of Martini 3–based CG systems, from simple molecules to polymers and large biomolecular complexes. In the following sections, we describe the architecture and implementation of the new Polymer Builder, the enhancements made to the Molecule and System Builders, and the expansion of the MAD:Database. We also present a set of case studies that demonstrate the new capabilities of the platform in constructing polymers, oligomeric proteins assemblies, and other systems. Finally, we discuss current limitations, future directions for development, and the broader role of MAD in the evolving Martini ecosystem.

## Material and Methods

### Overall Organization and Philosophy

The MAD Server was designed as a public repository of CG molecules integrated with a suite of CG tools. Its backend infrastructure provides storage and computational resources, enabling users to assemble and generate complete molecular CG systems ready for MD simulations.

The new version of MAD expands both the available tools and molecular resources while enhancing interoperability between tools and CG model data, forming a true MAD:Ecosystem (Figure 1). In most cases, CG models produced by one tool can be directly used as input for another. The MAD:Database anchors this ecosystem, with most deposited models accessible across multiple MAD tools.

**Figure 1:**
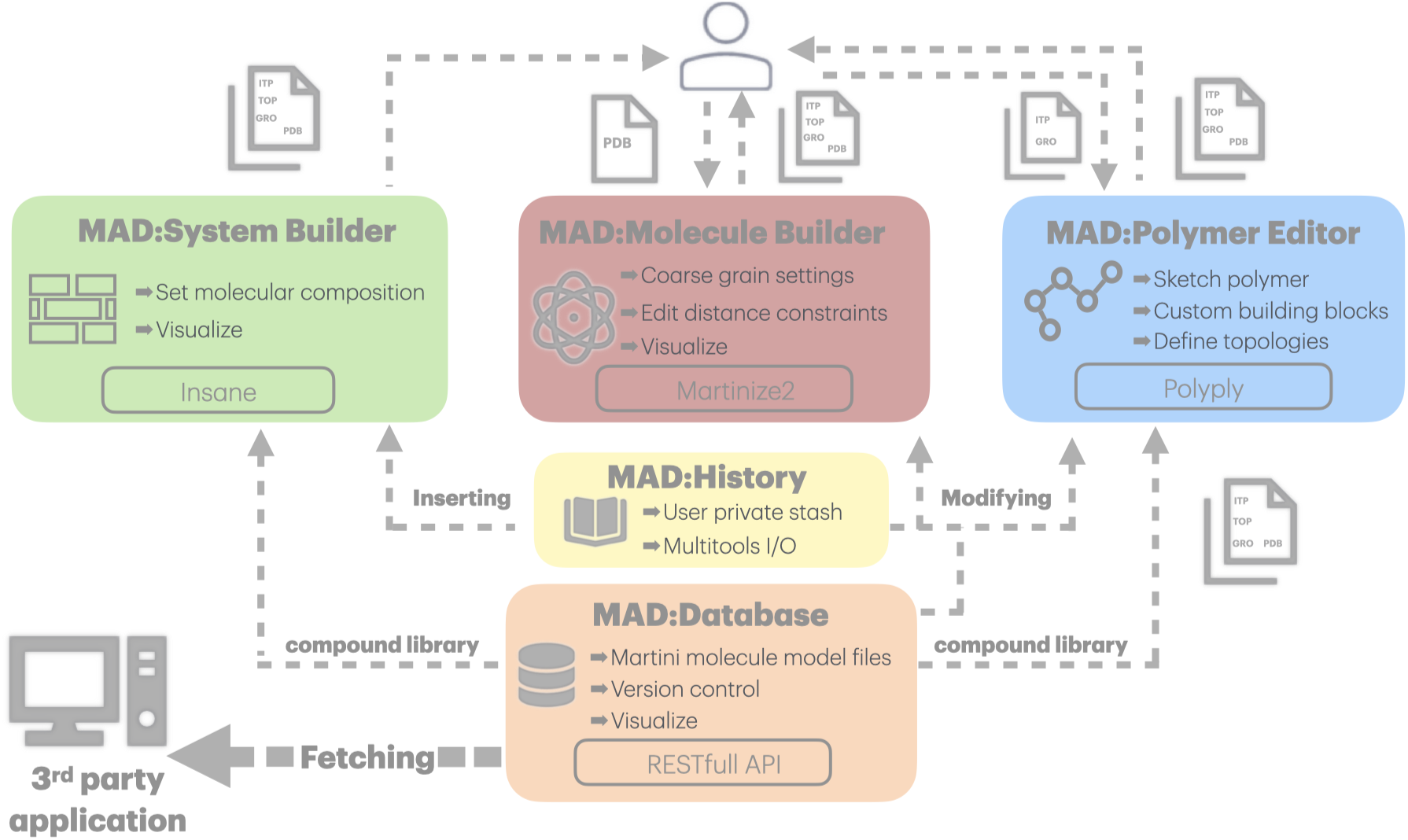
The MAD server workflow was expanded with the addition of the MAD:Polymer Editor. MAD users are granted a MAD:History private storage which backs up all their models generated by MAD:Polymer Editor and MAD:Molecule Builder. The new MAD API offers programmatic access to all MAD:Database content. The compounds stored in the MAD:Database can be imported in the MAD:Polymer Editor and the MAD:System Builder.

### The Polymer Editor

The implementation of the MAD:Polymer Editor (https://mad.ens-lyon.fr/polymer) is the latest major extension to the new MAD webserver. It functions as a graphical user interface and workflow manager for the polyply^8^ package. Through polyply, it assists users in generating input files and initial system coordinates for simulating disordered (bio)macromolecules such as synthetic polymers, polypeptides, or polysaccharides.

The MAD:Polymer Editor represents polymer molecules as graphs. The nodes of the graph represent building blocks, such as single beads, residues, or larger fragments like proteins. The edges follow the molecule’s covalent connectivity. From this graph, the simulation input files are generated by combining predefined building blocks specifying bead types and bonded interactions, along with links that modify these interactions when an edge is present. In the graphical interface (Figure 2), available building blocks can be selected and connected via the left-hand multipurpose menu (Figure 2A). The resulting graph is then displayed in the right-hand-side molecule sketcher section (Figure 2B).

**Figure 2:**
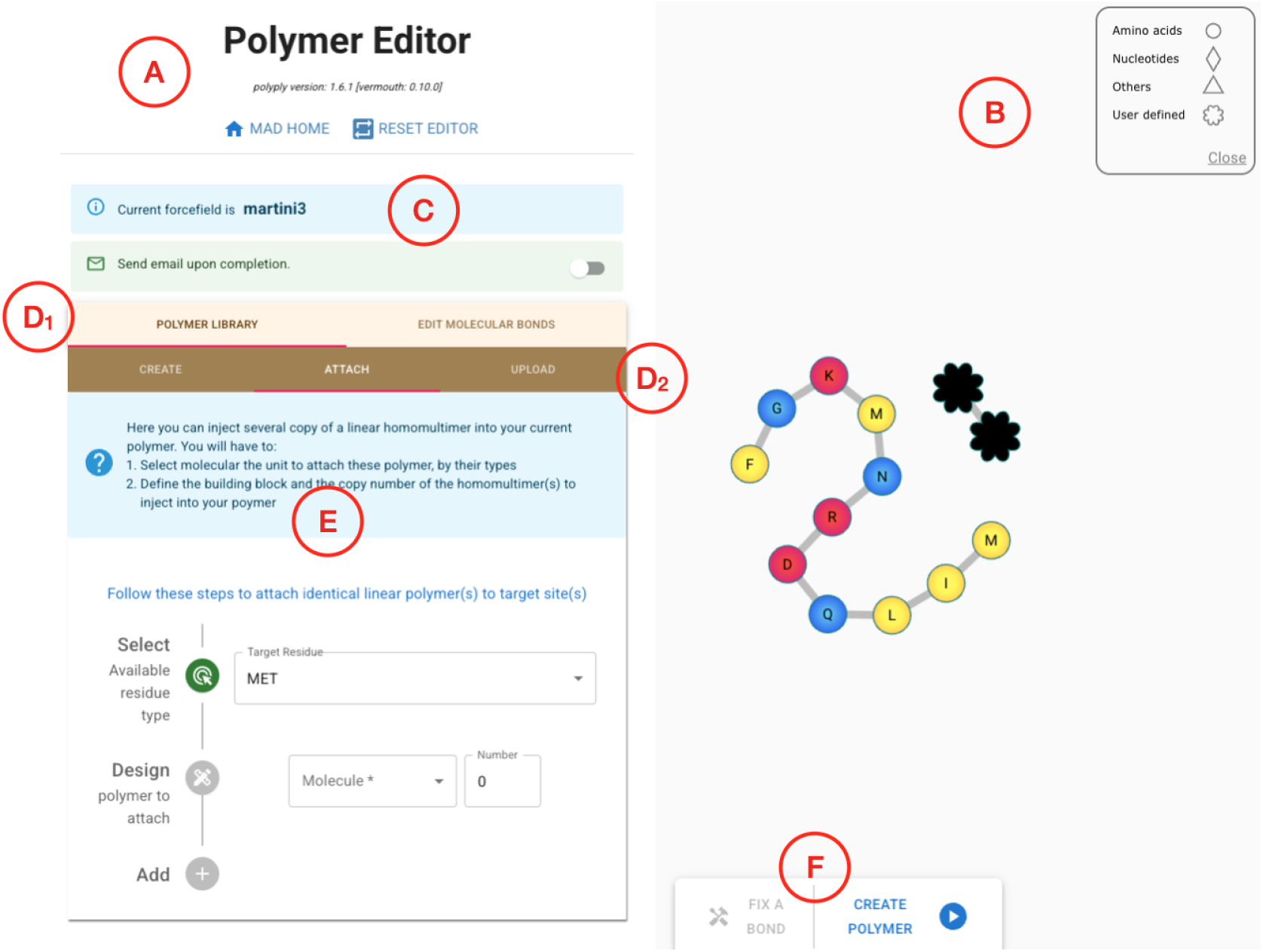
General layout of the MAD:Polymer Editor user interface. A) left-end menu panel with library versions on top, B) right-end interactive polymer sketcher. The panel section features: C) the force field currently selected, D1) Top panel section with tab menus. The top menus: Polymer Library menus (currently selected) and the molecular bond edition give access to specific action menus (D2) such as the showcased “Attach Polymer” which allows for the addition of several identical polymers branches at specific positions in the currently sketched molecule. The bottom console (F) supports the processing of the molecule and provides easy access to fix improper molecular bond.

The MAD:Polymer Editor allows users to select building blocks from the library shipped with the polyply package expanded to all molecules registered in the MAD:Database (Figure 2D1). Thus, peptides and lipid design can benefit respectively from the latest IDP extension^14^ and the latest lipid tails models.^20^ Single building blocks can be added to the sketch one by one or as linearly connected copies via the designated menu section (Figure 2E). Multiple building blocks can be attached to several target sites simultaneously. Additionally, users can specify the building block graph, or parts thereof, by uploading GROMACS ITP files or polyply JSON files. Linear peptides can be provided in FASTA format. Once added to the MAD:Polymer Editor, all building blocks and complex polymers remain accessible through the Polymer library menu (Figure 2D1) for the duration of the user’s session.

The molecule sketcher section allows users to dynamically interact with the graph representation of the polymer. Each node can be labeled with a unique or multiple residue identifiers, the former ones are presented by circular or polygonal shapes, the latter by a flower-like shape (Figure 2B). The molecule sketcher supports copying and pasting arbitrary graph components and creating or deleting groups of nodes and links on the fly. The nodes also support a right-click contextual menu to perform specific remove, copy or connect actions. This feature facilitates the construction of large, complex graphs representing molecules composed of many different monomers and fragments.

Once sketched, the polymer can be submitted to the MAD server through the MAD:Polymer Editor console (Figure 2F) for processing. The server first attempts to generate the polymer ITP parameter file in a fail-safe environment using the polyply gen params routine. This step may fail if bond parameters between polymer elements are unknown to the MAD:Polymer Editor (eg: missing uploaded files). In such cases, the MAD server captures the exceptions and reports them to the MAD:Polymer Editor interface, where erroneous links are highlighted in red within the polymer sketch. For each highlighted link, a dedicated window presents possible bead types and bond parameters for user selection, with sensible defaults pre-assigned. These user-defined links are intended as placeholders, allowing users to later refine and improve the bonded parameters during subsequent parametrization or simulation setup steps outside the MAD interface.

Once all erroneous links are resolved—or if none are detected—the server proceeds to generate the polymer’s starting coordinates, perform energy minimization, and produce a relaxed final structure. Upon completion, the MAD:Polymer Editor displays a “ball and stick” representation of the resulting polymer along with download links for ITP, TOP, GRO, and PDB files. The user may then store the generated structure in their personal MAD:History.

The MAD:Polymer Editor can start from a previously provided molecular structure or from the user’s personal MAD:History.

### Updated Database and API

The MAD:Database was originally designed to store a broad range of CG molecules and has grown, through user contributions, to include 1,274 CG models for 976 molecules, organized into the following categories: carbohydrates, polymers, amino acids, lipids, ions, phytochemicals, solvents, surfactants, synthetic nanoparticles, and small molecules. The recent addition of the latest Martini 3 lipidome set^20^ has greatly increased the model numbers (Table 1).

**Table 1:**
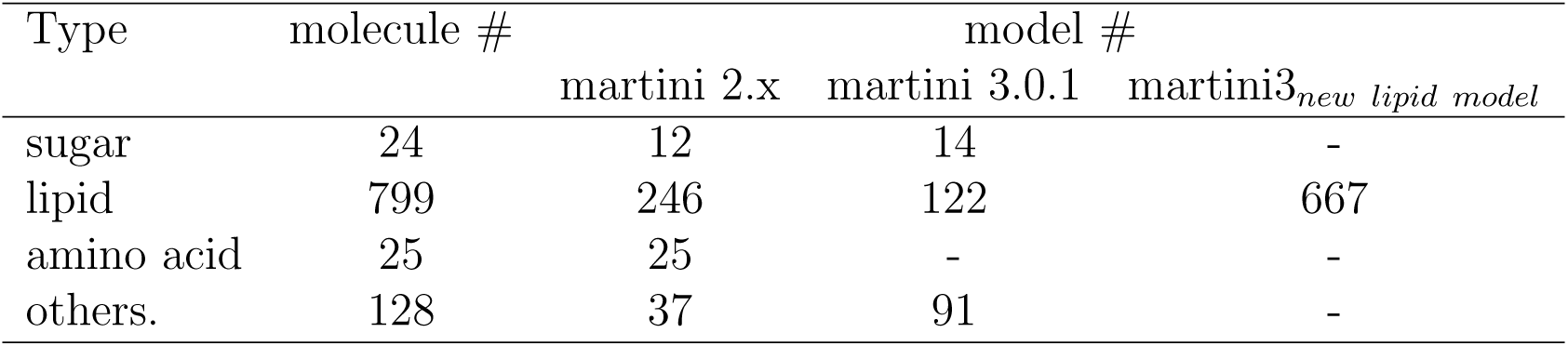
MAD:Database content overview.

| Type | molecule # | model # |  |  |
| --- | --- | --- | --- | --- |
|  |  | martini 2.x | martini 3.0.1 | martini3<sub>new lipid model</sub> |
| sugar | 24 | 12 | 14 | - |
| lipid | 799 | 246 | 122 | 667 |
| amino acid | 25 | 25 | - | - |
| others. | 128 | 37 | 91 | - |

The key functional improvement of the MAD:Database is its newly established interoperability with the MAD:Ecosystem. In particular, the MAD:System Builder and the MAD:Polymer Editor can now automatically retrieve, use, and combine any compatible CG models directly from the hundreds of entries in the MAD:Database. This greatly expands the original molecular coverage of these tools and allows users to reuse products from their MAD:History across other MAD:Ecosystem tools.

The MAD:Database is now accessible via a REST API, enabling programmatic interaction. This API offers fast access to MAD:Database data and resources, supports batch data processing, allows for task automation, and facilitates integration with external tools and systems. As a result, third-party developers can leverage the API to build new applications that utilize the data and capabilities of the MAD:Database.

The API protocol implements **GET** and **LIST** actions. The **GET** action resource url has the following format https://mad.ens-lyon.fr/api/molecule/get/[force_field]/[molecule_name][extension]/[version], where the url parameters are described in Table 2. The elements returned by a **GET** resource are CG model files.

**Table 2:** GET parameters of the MAD REST API.

| parameter | mandatory | default | description |
| --- | --- | --- | --- |
| force_field | yes | None | force field version (e.g. martini3001) |
| molecule_name | no | None | alias of molecule (e.g. POPE) |
| extension | no | .zip | bundle or specific file (.itp, .gro, .pdb) |
| version | no | 1.0 | model version number |

The **LIST** action resource url has the following format https://mad.ens-lyon.fr/api/molecule/list?[parameter]=[value], where parameter can be categories, name, force field or alias. Multiple parameters can be provided (e.g. http://mad.ens-lyon.fr/api/molecule/list?categories=Sugars&name=Sucrose). The element returned by a **LIST** resource is a JSON file listing MAD:Database entries matching the request. The document is limited to 200 entries, but the url parameters *skip* and *limit* allow for extended fetching.

### Improvements of existing tools and resources

The MAD:Molecule Builder can process an all-atom structure provided by the user to generate the corresponding CG structure and topology. It is built on top of the martinize2 program^10^ and has been upgraded to leverage recent developments in this software (v0.15.0) and in the Martini Protein Model.^14,15^

CG models that omit explicit hydrogen bond terms often require secondary and tertiary structural bias to preserve protein folding during simulations. Previously, the Molecule Builder supported structural bias exclusively through elastic networks and was limited to monomeric structures. With the recent implementation of GōMartini 3 in martinize2,^15^ the MAD:Molecule Builder now allows interactive configuration of Gō-bond Lennard-Jones potentials for any oligomeric state. Gō-bond networks can thus be defined both within and across molecular subunits. When editing groups of Gō restraints simultaneously (e.g., interchain restraints), their Lennard-Jones *ε* parameters can be adjusted directly through the interface. In addition, specific water–protein interaction biases can now be set independently for *α*-helix, *β*-strand, and coil regions.

With the recent release of the Martini3-IDP extension^14^ for disordered protein regions, MAD:Molecule Builder has also been equipped with an interactive widget to guide the user in the definition of protein subsequences to be modeled as disordered.

The MAD:System Builder has been upgraded to accommodate the growing number of lipid CG models. It can now build membranes using Martini 2, Martini 3, and the latest Martini 3 lipidome models.^20^ The pool of available models has expanded, as all entries from the Lipids section of the MAD:Database can now be imported directly. The system also alerts users when incompatible models are combined.

### Case studies

To illustrate the latest development of the MAD:Polymer Editor and the overall improvements of the MAD:Ecosystem, we present different test cases.

#### Construction of synthetic and biological disordered polymers

The capabilities and flexibility of the MAD:Polymer Editor, can be illustrated using a set of examples comprised of polysaccharides, intrinsically disordered proteins (IDP), and synthetic polymers with increasing architectural complexity. Macromolecular structures were built using the interactive graph-based MAD:Polymer Editor, with some examples combining building blocks originating from polyply and retrieved from the MAD:Database. The resulting models were exported as standard GROMACS input files and tested for compatibility with Martini 3 simulation protocols.

A dextran polymer was constructed by repeating a glucose-based monomer from the polyply library of the MAD:Polymer Editor (Figure 3A). Dextran is a polysaccharide composed of *α*-1,6–linked glucose units and is well known for its high water solubility compared to other polyglucoses. While high–molecular weight dextrans are typically branched, lower degrees of polymerization are often only weakly branched or effectively linear. Accordingly, the dextran model generated here corresponds to a linear polysaccharide chain. Dextran has previously served as a benchmark system for CG simulations of polysaccharides during the development of the Martini 3 carbohydrate model,^12^ where its solution properties behavior were shown to be well captured. This example illustrates the straightforward construction of carbohydrate-based polymers using predefined building blocks within the MAD:Polymer Editor.

**Figure 3:**
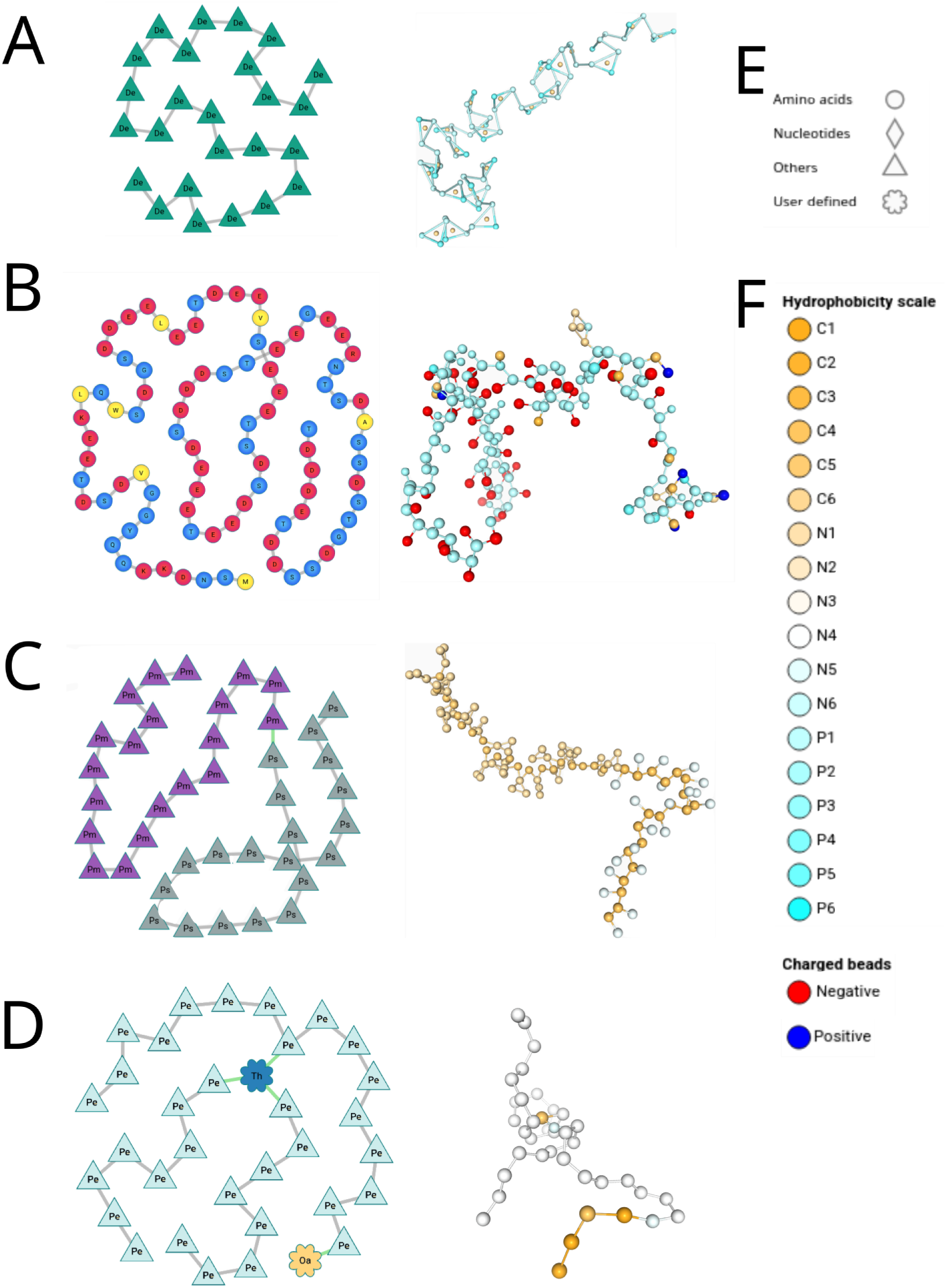
Case studies of synthetic and biological disordered polymers constructed with the MAD:Polymer Editor. Panels A–D show, for each system, the corresponding graph representation used during construction (left) and the resulting CG structure (right). In the graph representations, colors are used solely to differentiate monomer types; missing bonds originating from polyply definitions are indicated in green and were subsequently resolved interactively within the MAD:Polymer Editor. Colors in the CG structures follow the bead-type color scale defined in panel F. (A) Linear dextran polymer composed of 20 glucose monomers. (B) Intrinsically disordered protein corresponding to residues 1–90 of the Sic1 protein from *Saccharomyces cerevisiae*. (C) Amphiphilic block copolymer composed of polystyrene and polymethyl acrylate blocks (PS_20_–PMA_20_). (D) TWEEN-like PEG–lipid architecture composed of branched PEG chains (PEG_8_) connected to a hydrophobic moiety (tetrahydrofuran) and oleic acid. (E) Legend for the graph representations. (F) Legend for the CG structures.

The same workflow was then applied to the generation of a sequence-defined disordered protein, as example of a biological polymer. More specifically, we chose the intrinsically disordered cyclin-dependent kinase inhibitor Sic1 from *Saccharomyces cerevisiae*, a prototypical IDP that has been extensively studied experimentally and computationally.^14,22,23^ A truncated construct corresponding to residues 1-90 was generated directly from a FASTA sequence using the MAD: Polymer Editor (Figure 3B), leveraging the Martini3-IDP^14^ model without requiring a predefined atomistic structure. This example highlights how the MAD:Polymer Editor can be used for sequence-driven, intrinsically disordered biomolecular systems.

To demonstrate the assembly of heterogeneous synthetic polymer architectures, an amphiphilic block copolymer composed of polystyrene (PS) and polymethyl acrylate (PMA) was constructed with the same graph-based approach used for dextran (Figure 3C). Building blocks corresponding to each polymer segment with 20 monomers were selected from the MAD:Polymer Editor and connected interactively. At the junction between the two polymer blocks, a missing bond was detected and resolved using the bond editing tools provided by the MAD:Polymer Editor, illustrating how manual addition of bonds can be performed in polymer systems. The resulting CG models demonstrate how chemically distinct polymer blocks of user-defined lengths can be combined within a single workflow and exported as simulation-ready topologies compatible with the Martini 3 force field.

Finally, a branched polyethylene glycol (PEG)-based lipid was constructed to illustrate cases where nontrivial connectivity and building blocks leads to missing parameters (Figure 3D). The architecture was inspired by PEG–lipid surfactants such as polysorbates (e.g., TWEEN-like molecules), which combine branched PEG chains with hydrophobic moieties and are commonly used in formulation and nanocarrier contexts.^24,25^ In this example, a tetrahydrofuran moiety and oleic acid were taken from the MAD:Database and incorporated into the PEG-branched architecture with MAD:Polymer Editor. During topology generation, undefined bonds involving the tetrahydrofuran group, the PEG branches, and their connection to a oleic acid were automatically detected and highlighted within the MAD:Polymer Editor interface. These connections were resolved interactively using the bond editing tools, with default parameters provided as placeholders. Bead chemical type of fatty acid head was changed to a SN4a considering the connectivity with the PEG moiety. This example illustrate how polymer models generated with the MAD:Polymer Editor can be combined with molecules of MAD:Database.

For all examples, the generated macromolecular models could be visualized directly within the MAD web interface of MAD:Molecule Builder and were successfully exported as complete topology and coordinate files. The systems were also subsequently transferred to the MAD:System Builder and solvated in water boxes for benchmarking purposes. Short CG simulations confirmed that the resulting systems are stable and suitable for use in standard Martini 3 simulation workflows.

#### Tuning the Gō-bond network of large macro-molecular complexes

The MAD Server can also facilitate the modeling of large macro-molecular complexes, an otherwise daunting task. To illustrate a typical workflow, we modeled the Enoyl-acyl-carrier protein reductase (InhA) from Mycobacterium tuberculosis (MTB). It is essential for the survival of this this pathogen and is the primary target of isoniazid, one of the first-line drugs used for tuberculosis treatment. InhA is therefore a classical drug target and has been widely studied over the years.^26^ Missense mutations in the coding region of this protein have been associated with isoniazid resistance; however, the underlying resistance mechanisms remain poorly understood. CG MD simulations offer a powerful approach to study this system.^27,28^

MAD:Molecule Builder was used to generate monomeric and tetrameric CG systems for wild-type and mutant InhA using Martini 3 force field in a dynamic and visual workflow. In a preliminary stage, protein models (unpublished structures) were stripped of water molecules and ligands to create clean PDB files. Cleaned protein models were uploaded to the MAD:Molecule Builder using the generate molecule option, with virtual Gō Sites as Mode, and both Side-Chain fix and Cysteine Bridges enabled (Figure 4A). MAD correctly constructed the CG model (Figure 4B). Using the EDIT option in the Virtual Gō Bonds section, both intra- and inter-chain virtual Gō bonds could be inspected (Figure 4C).

**Figure 4:**
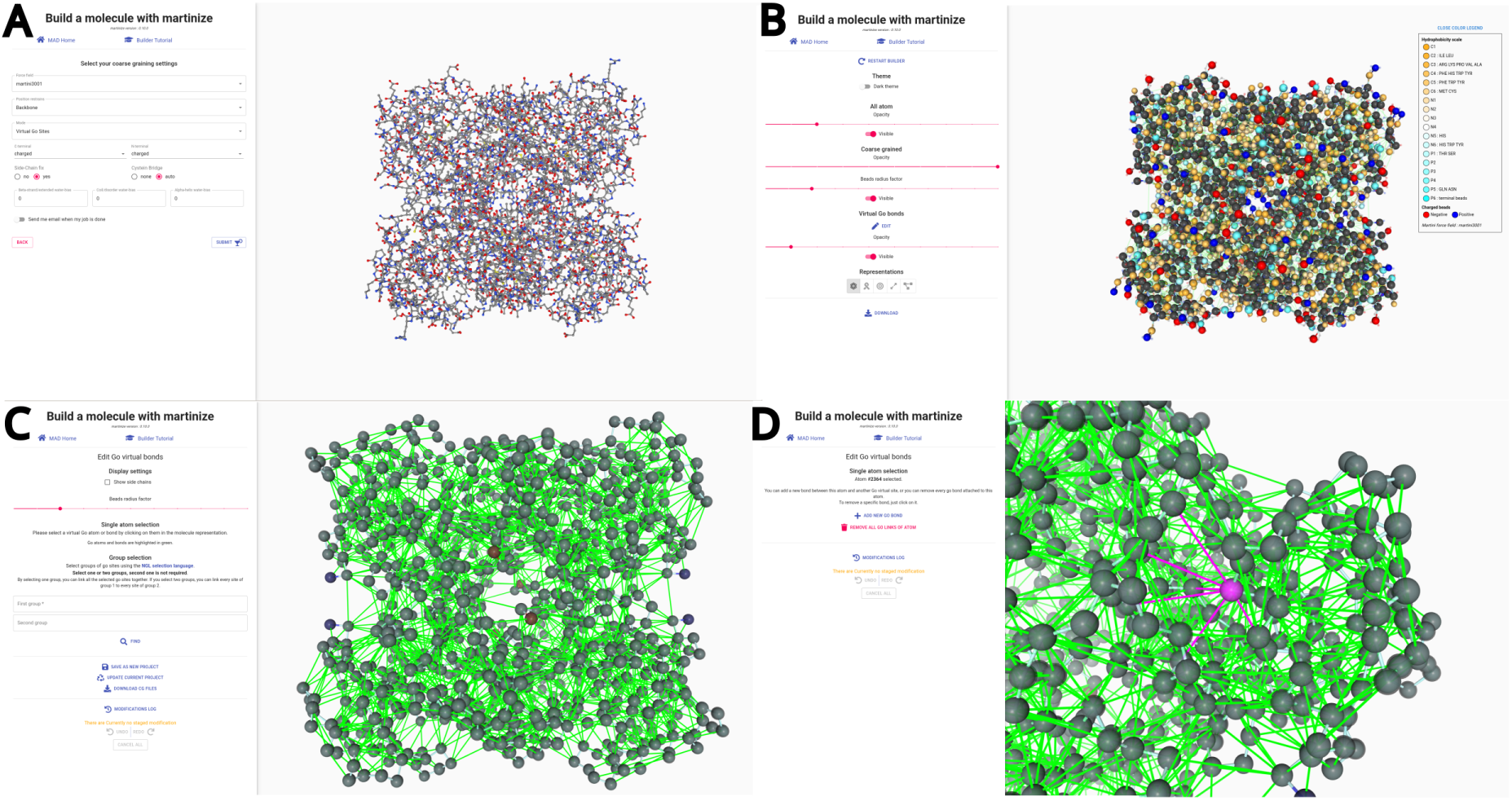
Building and editing a large CG model with Gō bonds in MAD:Molecule Builder. A - Initial upload of all atom PDB models and parameters used for CG model construction. B - Coarse Grained model. C - Virtual Gō bonds shown as green lines and beads as grey spheres. D - Closer look at residue specific Gō bonds network, highlighted in pink and easily editable.

The MAD:Molecule Builder automatically defines Gō bonds from the martinize2-generated contact map, retaining only contacts whose Martini backbone beads pairwise distances fall within the default cutoff criteria.^15^ Because the reference structure represents a single conformation, the resulting Gō network may impose overly restrictive constraints, limiting natural protein dynamics in MD simulations. Conversely, additional Gō contacts not present in the reference structure may be required to facilitate functional motions during simulation. In the default CG model of InhA WT generated by MAD:Molecule Builder, residues 164 and 165 of each protomer were assigned multiple interchain Gō bonds. These interactions reduced physiologically relevant chain fluctuations in subsequent MD test simulations when compared with reference all-atom simulations. To address this problem, the model was interactively modified within the MAD:Molecule Builder by removing the corresponding Gō bonds using the editor ((Figure 4D). The resulting CG model was then used in long MD simulations to investigate the effects of missense mutations in the Mycobacterium tuberculosis InhA coding region. To our knowledge, MAD:Molecule Builder is currently the only coarse-graining tool that enables interactive editing of inter- or intra-chain Gō bonds.

#### Modeling partially disordered protein

To showcase the ability of the MAD:Molecule Builder at modeling disordered protein domains, it was used to build a Martini 3 model of the membrane-bound monomeric *α*-synuclein (*α*S). *α*S is a neuronal protein implicated in synaptic vesicle trafficking and Parkinson’s disease, and represents a prototypical IDP capable of functional folding upon membrane interaction. *α*S is remarkable for its ability to adopt a diverse range of conformational states.^29^ Under physiological conditions, the *α*S monomer is largely disordered in the cytosol but adopts a highly ordered *α*S-helical conformation upon membrane binding. The N-terminal segment (residues 1–95) contains a repeating KTKEGV motif that mediates lipid membrane binding. Upon adsorption to the membrane, this segment folds into an amphipathic *α*-helix, as demonstrated by biophysical studies.^29,30^ Experimental evidence further indicates that the helix lies parallel to the membrane interface and inserts shallowly into the bilayer, with its center located in the upper acyl region just below the phosphate groups.^29,31^ The remaining C–terminal acidic region does not interact with the membrane and remains unfolded.

For this model, we used as reference an atomistic structure of micelle-bound *α*S obtained by NMR^30^ (PDB ID: 1XQ8^32^). In this structure, residues 3–92 adopt a helical conformation, with residues 38–44 forming a short flexible linker between two helices, while the C-terminal tail remains disordered. To construct the Martini 3 model of *α*S, we adopted a two-pronged approach. The *α*-helical regions were represented using the regular Martini 3 model combined with GōMartini to preserve secondary structure throughout the simulation. In addition, GōMartini was employed to introduce a water–protein interaction bias, stabilizing the association of the helices with the membrane. The intrinsically disordered C-terminal region was modeled using Martini 3-IDP extension,^14^ which provides an improved description of disordered chain conformations.

To generate this hybrid model with MAD:Molecule Builder, we selected “martini3IDP” as the force field (Figure 5A). In practice, this is identical to the standard Martini 3 force field, except for the regions explicitly defined as disordered using the “Define disordered region” option (Figure 5B). In these regions, optimized bonded potentials for intrinsically disordered residues are applied, and any secondary or tertiary structure biases—such as elastic networks or GōMartini restraints—are automatically removed. We then chose GōMartini as the structural bias (“Mode: Virtual Gō sites”) to maintain the *α*-helical segments, and apply a –1 kJ/mol *α*-helical water bias to fine-tune protein–membrane interactions (Figure 5A). After submitting the job, MAD:Molecule Builder provided the option to refine the GōMartini network before generating the final CG topology. In this step, we manually adjusted the Gō bonds to tune the flexibility of the model. Specifically, we removed the Gō interactions constraining the linker region between the two *α*-helices, thereby allowing each helix to move independently while maintaining the overall folded character of the helical segments (Figure 5C,D). At this point, the Martini 3 *α*S model was complete (Figure 5E). This example demonstrates how MAD:Molecule Builder facilitates the flexible and intuitive construction of hybrid Martini 3 models, seamlessly integrating structured and intrinsically disordered regions within the same protein.

**Figure 5:**
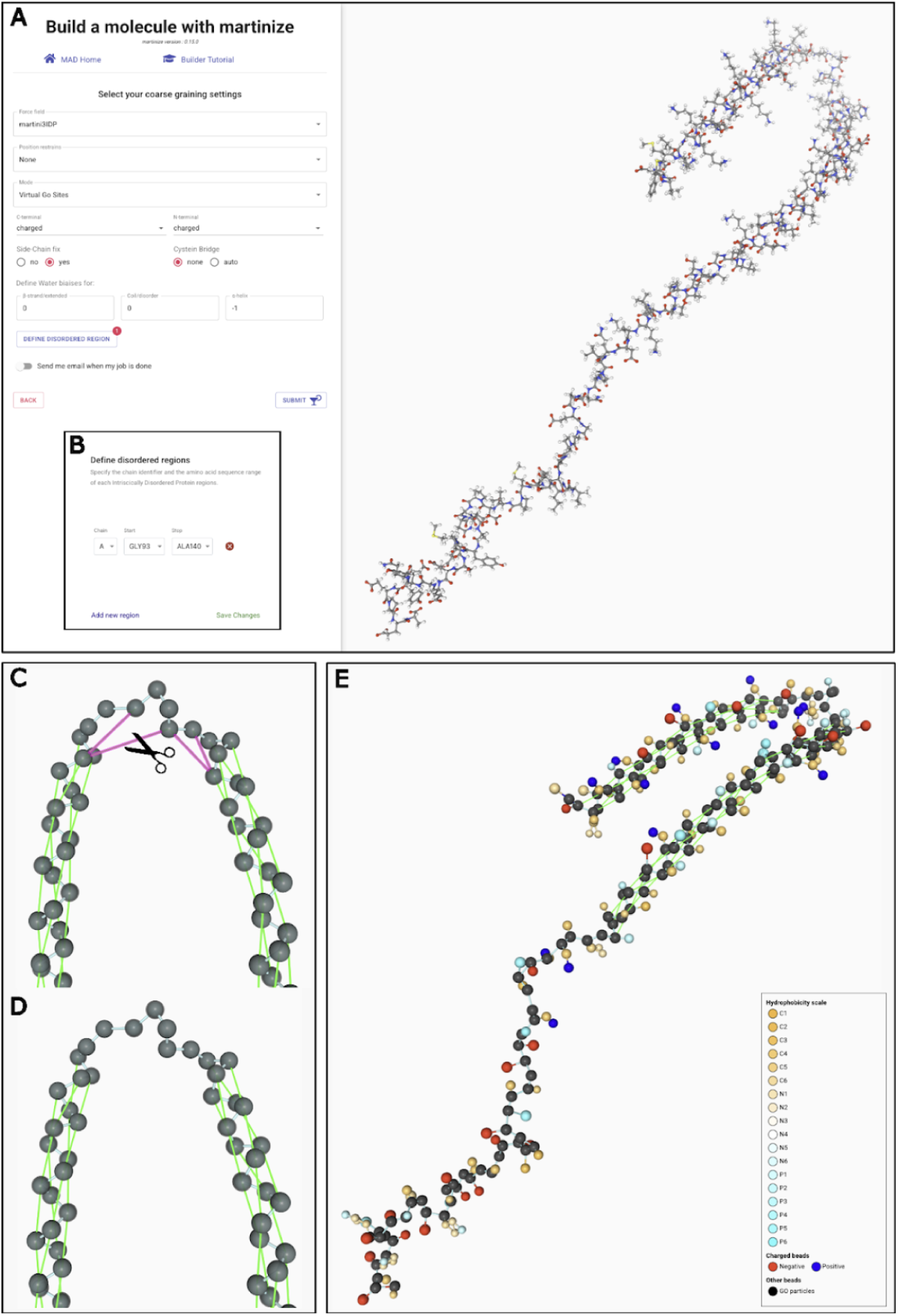
CG Modeling of a partially disordered protein with MAD:Molecule Builder. A) Initial upload of all atom PDB models and setting of the Martini 3-IDP force field extension. B) Selection of the disordered regions. C,D) Interactive editing of the Gō bonds. E) Display of the final model

## Conclusion and perspectives

This work presents a major update of the MAD web server, reflecting both the continued evolution of the Martini ecosystem and the changing needs of its user community since the release of the previous MAD version. ^9^ While earlier iterations of MAD primarily focused on providing access to Martini-ready compounds and basic system preparation workflows, the present release expands the scope of the platform and improves the integration between tools for CG model construction, with particular emphasis on compatibility with Martini 3.

A central addition of this update is the MAD Polymer Builder, which extends MAD beyond biomolecular systems and enables the construction of CG polymer models with diverse architectures and compositions. This functionality represents a clear expansion relative to the previous MAD release and complements existing polymer-oriented tools by providing a web-based environment tightly integrated with the MAD database. In cases where connectivity between monomers is missing from the library, the current implementation allows users to manually define bonded interactions, including bead selection, bead types and sizes, bond lengths, and force constants. While this manual editing workflow is functional, the visualization support during bond definition remains limited and could be improved to facilitate user interaction. Despite this flexibility, the lack of automated angle and dihedral generation remains a limitation. Future developments, including continued extensions of polyply^8^ and the incorporation of automated topology-generation strategies, potentially guided by analogy to existing database entries or by more advanced data-driven approaches, are expected to address these limitations. In this context, efforts toward automated CG parametrization, such as AutoMartini,^33,34^ PyCGTOOL^35^ and Bartender,^36^ highlight promising directions for reducing manual intervention and could inform future developments of automated workflows within MAD. The recent automated Martini 3 parametrization of disaccharides^37^ illustrates the potential of these approaches for carbohydrate-based polymers in MAD.

The MAD:Molecule Builder has been further extended and now supports most commonly used Martini protein modeling variants, including elastic network models, GōMartini representations, and water-biased interaction schemes. Compared to the previous MAD version, these additions provide increased flexibility for controlling protein stability and interaction behavior. More recent approaches such as OLIVES,^38^ which provide an alternative framework for modeling protein flexibility in Martini, are not yet implemented in MAD. At the same time, practical challenges remain when dealing with experimental structures that contain missing residues, inconsistent atom naming, or incomplete atomistic information. Future improvements aimed at automated pre-processing of protein structures would help facilitate model preparation in such cases. In addition, while post-translational modifications such as lipidations^39^ are supported by martinize2, MAD may be limited to cases where these modifications are already present in the reference atomistic structure, and they cannot yet be introduced de novo through the MAD interface. The same limitation applies to other post-translational modifications, including glycosylations, phosphorylations, and related chemical modifications. Ongoing developments of CG models within the Martini framework suggest that their more systematic integration into MAD may become feasible in the near future.

At the system assembly level, MAD continues to provide streamlined workflows for constructing simulation-ready environments through the System Builder module, which now provides a more extensive lipid library in line with recent Martini model releases.^20,21^ As the size and diversity of the lipid library increase, further improvements in search capabilities and molecular visualization within the builder would be beneficial to facilitate system setup. An additional extension that could further support users would be the inclusion of predefined complex membrane compositions^40–43^ as selectable templates, enabling the rapid construction of biologically relevant lipid mixtures without requiring manual specification of individual components. Some other limitations are inherited from the underlying insane engine,^11^ particularly with respect to the construction of systems containing multiple embedded macromolecules or more complex membrane geometries. In addition, users may need to manually pre-orient proteins prior to membrane insertion, although this represents a preprocessing step that could be readily integrated into the workflow in the future. Planned integration of alternative engines such as COBY^5^ and TS2CG^6,7^ is expected to progressively expand the range of accessible system architectures, including curved membranes and more complex lipid-based compartments.

Beyond functional extensions, this release also reflects changes informed by operational experience with the previous MAD deployment. During the lifetime of the MAD service, periods of platform instability limited accessibility and highlighted limitations in the organization of the codebase and in the deployment model. In the present update, these issues have motivated a reorganization of the platform, with increased emphasis on modularity, clearer separation of components, and improved portability of the infrastructure. These efforts aim to facilitate smoother transitions in the event of future infrastructural modifications and to improve overall robustness and usability for the community.

Finally, this update includes a significant expansion of the MAD:Database and improved integration between the different components of the MAD:Ecosystem. These developments are in line with broader directions in the field toward more integrated and flexible platforms for the construction of complex molecular systems, as illustrated by recent advances in the CHARMM-GUI Multicomponent Assembler^44^ and other developments for complex system assembly, including virus^45^ and cell models.^46^ Continued development will focus on strengthening interoperability, both within MAD and through closer interaction with community resources such as the Martini Force Field Initiative webpage and git repositories via programmatic interfaces.

A remaining challenge is that many Martini topologies released in the literature are not systematically deposited in MAD or other organized repositories, limiting their long-term accessibility and reuse. Establishing community-driven best practices—potentially including the expectation that newly published Martini models be deposited in public databases such as MAD, in a manner analogous to established practices for structural data (i.e., the PDB)—would substantially improve reproducibility, visibility, and collective model development. In parallel, initiatives such as the Molecular Dynamics Data Bank (MDDB)^47^ provide a complementary infrastructure focused on the long-term storage, curation, and dissemination of MD trajectories. In this perspective, MAD naturally complements MDDB by focusing on the generation, standardization, and dissemination of Martini topologies and starting structures, together supporting a more complete and FAIR CG simulation ecosystem.^48^ Related cloud-based approaches for running molecular simulations^49^ further highlight complementary directions for improving accessibility. In this context, future extensions of the MAD:Database and its API could benefit from emerging standards for CG molecular representation, such as CGsmiles,^50^ which provide a resolution-independent description of CG mappings and facilitate indexing, interoperability, and high-throughput applications. Together, these developments outline a clear direction for the continued evolution of MAD, building on the foundations established by earlier versions while progressively extending its applicability and scope.

## Authors Information

## Contributions

P.C.T.S. and G.L. designed the project. R.M., C.H., and G.L. performed the software development, while M.V. and P.C.T.S. contributed to data acquisition. F.G., L.B.A., N.O.R., M.V., and P.C.T.S. carried out the case studies. P.C.T.S. and G.L. wrote the manuscript with contributions from F.G., L.B.A., M.V. and N.O.R.. All authors reviewed and approved the final version of the manuscript.

## Acknowledgement

We thank all external users for testing MAD during its development. M.V., L.B.A., N.O.R. G.L., and P.C.T.S. acknowledge the support of the PSMN (Pôle Scientifique de Modélisation Numérique) and the Centre Blaise Pascal’s IT test platform at ENS de Lyon (Lyon, France) for access to computational resources. The platform operates the SIDUS solution developed by Emmanuel Quemener.^51^ P.C.T.S. and L.B.A. acknowledge support from the French National Center for Scientific Research (CNRS). Further funding of P.C.T.S. came from a research collaboration with Sanofi. F.G. acknowledges funding from the Klaus Tschira Foundation (Independent PostDoc Fellowship). S.J.M. is supported by funding from the European Research Council (ERC) under grant agreement No. 101053661 (“COMP-O-CELL”).

## Data and Software Availability

## Supplementary Material

### Coarse-grained molecular dynamics simulations

CG MD simulations were performed to validate the systems generated with the updated MAD:Polymer Editor and MAD:Molecule Builder and to verify their compatibility with standard Martini 3 simulation protocols. All simulations were carried out using the Martini 3 force field^2^ and followed the recommended simulation settings for Martini-based CG models.^52,53^

Simulations were performed with GROMACS 2024.x and 2025.x,^54^ using a leap-frog integration scheme.^55^ Neighbor lists were updated using the Verlet neighbor search algorithm,^56^ and nonbonded interactions were treated with a Verlet cutoff scheme.^57^ A typical CG time step of 20 fs was used for production simulations. Nonbonded interactions were treated using a Lennard-Jones cutoff of 1.1 nm with a potential-shift scheme, while long-range electrostatics were handled using either a reaction-field approach.^58^ Periodic boundary conditions were applied in all three dimensions. Temperature and pressure were maintained using the velocity-rescaling thermostat^59^ and the Parrinello–Rahman barostat,^60^ respectively, with coupling constants consistent with standard Martini 3 practice. The purpose of these simulations was not to perform detailed quantitative validation, but rather to confirm that the generated systems are stable, free of numerical instabilities, and suitable for production Martini 3 simulations. All showcased structure input files and corresponding CG data output are publicly available at the MAD codebase repository (https://gitbio.ens-lyon.fr/LBMC/damm/mad/paper2026_si). Specific topology files can be alternatively obtained from MAD:Database(https://mad.ens-lyon.fr/explore) or the martini force field initiative website (https://github.com/Martini-Force-Field-Initiative).

